## Supplementary figures and images for "Infinite Physical Monkey: Do Deep Learning Methods Really Perform Better in Conformation Generation?"

### TOC

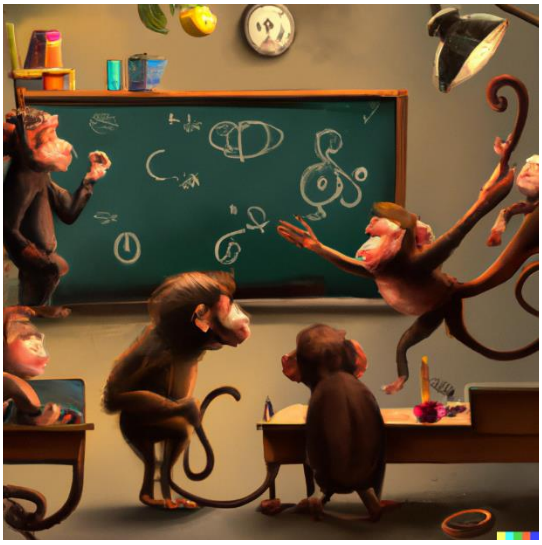
